# Fecal microbial transplant abates tolerance to methylone-induced hyperthermia

**DOI:** 10.1101/2021.01.11.426194

**Authors:** Robert Goldsmith, Amal Aburahma, Sudhan Pachhain, Sayantan Roy Choudhury, Vipa Phuntumart, Ray Larsen, Jon E. Sprague

## Abstract

The microbiome-gut-brain axis has been implicated in multiple bodily systems and pathologies, and intentional manipulation of the gut-microbiome has yielded clinically significant results. Here, we examined the effects of bi-directional fecal microbial transplants (FMT) between methylone-induced hyperthermic tolerant (MHT) and methylone-naïve (MN) rats. Rats treated with methylone once per week developed tolerance to methylone-induced hyperthermia by the fourth week. Once tolerant, daily bi-directional FMT between the two groups were performed for seven days prior to the next methylone treatment. The FMT abated the developed tolerance in the MHT group. When treated with methylone for the first time following FMT, recipient MN rats displayed significant tolerance to hyperthermia despite it being their initial drug treatment. Post-FMT, MHT rats displayed elevations in norepinephrine and expression of *UCP1*, *UCP3* and *TGR5* in brown adipose tissue, with reductions in expression of *TGR5* and *UCP3* in skeletal muscle. The pre- and post-FMT methylone tolerance phenotypes of transplant recipients are concurrent with changes in the relative abundance of several Classes of *Proteobacteria*, most evident for *Gammaproteobacter* and *Alphaproteobacter.* MHT recipients demonstrated a marked increase in the relative proportion of the *Firmicutes* Class *Erysipelotrichia*. These findings suggest that transplantation of gut-microbiomes can confer phenotypic responses to a drug.

## Introduction

Use of the sympathomimetic agent methylone, the β-keto analog of 3,4-methylenedioxy-methamphetamine (MDMA), in warm ambient environments has been shown to induce an acute rise in body temperature [1], and fatal hyperthermia upon ingestion of methylone has been documented in case reports [2]. Hyperthermia mediated by sympathomimetic agents, such as MDMA and methylone, has both central and peripheral mechanisms. Centrally, activation of dopaminergic [3] and serotonergic [4] receptors in thermoregulatory circuits in the hypothalamus [5,6,7] contribute to the activation of the sympathetic nervous system (SNS). Peripherally, elevated levels of circulating norepinephrine acting at the α_1_-adrenergic receptor induce peripheral vasoconstriction, resulting in a decrease of heat dissipation [8]. Concurrently, activity of norepinephrine at the β_3_-adrenergic receptor results in increases of free fatty acids and the activation of mitochondrial uncoupling proteins (UCP). These UCP enzymes then facilitate proton flux from the intermembrane space to the mitochondrial matrix in a manner that sheds the kinetic energy as heat instead of producing ATP, otherwise known as non-shivering thermogenesis [9]. Despite the neural pathways being well-documented, there is preliminary evidence of another contributor to the activation of sympathomimetic-induced thermogenesis: the microbiome [10].

Recent studies have demonstrated that the microbiome-gut-brain axis plays a major role in maintaining body temperature regulation [11, 12]. Li et al. [12] demonstrated that gut microbiome depletion as a result of treatment with multiple cocktails of antibiotics altered thermogenesis in mice exposed to cold environmental temperatures. The antibiotic treated mice displayed significantly lower core body temperatures at ambient room temperature, consistent with decreased *UCP1* gene expression in brown adipose tissue. Ridge et al. [10] demonstrated that the administration of antibiotics for 14 days prior to MDMA treatment significantly reduced core body temperature increases, and blunted expression of *UCP1*, *UCP3* and the bile acid receptor gene *TGR5* (a regulator of UCP expression). These latter findings suggest a potential link between the microbiome-gut-brain axis and sympathomimetic-induced hyperthermia.

In addition to thermogenesis, the microbiome-gut-brain axis has become increasingly implicated in a variety of physiologies and conditions, including obesity, cancer, and neurodegenerative disorders such as Parkinson’s Disease and Alzheimer’s [13, 14, 15], and has quickly become a major target area in attempts to advance our understanding of the bacterial influence on health and disease. Naturally, researchers have begun to manipulate the microbiome in attempts to modulate these pathways and have begun to elucidate the communication between the gut, the brain, endocrine, and immune systems. To date, microbial manipulations have shown to effectively mitigate depressive symptoms in mice, demonstrating the ability to modulate mood through selective transfer of the microbiome [16]. In other medicinal areas, fecal microbial transplants (FMT) have been used to successfully treat inflammatory bowel disease [17] and *Clostridium difficile* infection [18], further establishing that the intentional manipulation of the gut microbiome can result in novel and clinically relevant outcomes. While previous experiments have indicated that microbial manipulations are capable of treating and inducing disease states, to date no analyses have been performed to explore the role they play in development of phenotypic responses such as drug-induced hyperthermia.

In the present study, we hypothesized that chronic exposure to methylone might drive changes in bacterial populations within the gastrointestinal tract that could contribute to the methylone hyperthermic tolerance development [19] shown in our previous work. If so, then transferring the microbiomes of methylone-induced hyperthermic tolerant rats to animals naïve to methylone could replicate the tolerance effect in the absence of chronic drug exposure. Furthermore, we sought to discover whether the bi-directional transfer of methylone-naïve rat microbiomes to the tolerant rats would be sufficient to eliminate their developed hyperthermia tolerance to chronic methylone treatment.

## Materials and methods

### Animals

Adult, male (N=12, 275-300 g) Sprague-Dawley (*Rattus norvegicus domesticus*) rats were obtained from Envigo (Indianapolis, IN). Animals were housed one per cage (cage size: 21.0 × 41.9 × 20.3 cm) and maintained on a 12:12 h light/dark schedule. To maximize the thermogenic response, animals were maintained at an ambient temperature of 25°C to 28°C and fed a minimum 10% fat diet [20,21]. Animals received food in the form of a ground powder in glass container in order to habituate them for fecal microbial transplant methods. Animal maintenance and research were conducted in accordance with the eighth edition of the Guide for the Care and Use of Laboratory Animals; as adopted and promulgated by the National Institutes of Health, with protocols approved by the Bowling Green State University Animal Care and Use Committee.

#### Drug and Chemicals

Racemic methylone was obtained from Cayman Chemicals (Ann Arbor, MI) as a hydrochloride salt. On the day of the study, methylone solutions were made fresh at a concentration of 10 mg/mL in 0.9% normal saline. All other chemicals and reagents were obtained from Sigma Chemical (St. Louis, MO).

### Induction of Hyperthermia Tolerance

Male rats were randomly assigned into two groups of six (6) each, the first group being the treatment group and the second serving as the saline controls. On testing day, all subjects were weighed prior to drug challenge, and a core temperature reading was taken with a rectal thermometer at time zero. On treatment days, the ambient temperature averaged 27.4 ± 0.12°C. Following the first temperature measurement each week, the male treatment group received a 10 mg/kg subcutaneous (sc) dose of methylone, and the control group received an equal volume of saline solution (sc). Following drug challenge, core temperature readings were recorded at the 30-, 60- and 90-minute time points. This treatment schedule was maintained once a week for a total of four consecutive weeks, until the hyperthermic response of the methylone treatment group was statistically insignificant. Those animals treated weekly for four weeks with methylone were designated as methylone hyperthermic-tolerant (MHT) and those that received only saline for four weeks were designated methylone-naïve (MN).

### Fecal Microbial Transplant

As mentioned previously, animals had up to this point become habituated to consuming ground food powder. To initiate fecal microbial transplant, cage fecal droppings from the weekly period directly prior to the hyperthermia-tolerant treatment were collected from each cage and pooled by MHT and MN groups. Droppings were ground into powder in a mortar and pestle and mixed in with food powder in a 15% (w/w) ratio according to previous literature [22]. Beginning the same day after the treatment in which methylone rats did not exhibit statistically significant hyperthermia, bi-directional FMT was conducted where each rat received ab libitum access to FMT food powder of alternate group. This feeding schedule continued every day for 7 days until the next methylone treatment schedule. Food consumption was monitored daily.

Upon final treatment day (seven days from previous methylone dose during which FMT was administered daily), all 12 rats received methylone treatment at 10 mg/kg and temperature measurements were recorded as explained previously. After 90-minutes, rats were euthanized with CO_2_ and cardiac punctures were performed to collect blood samples. Plasma samples were stored at −20°C. BAT and SKM, namely the gastrocnemius, were removed and flash frozen with liquid nitrogen, before being stored at −80°C.

### RNA isolation and qRT-PCR

Total RNA was purified from homogenized SKM and BAT tissues using PureZOL^™^ RNA Isolation reagent (BioRad, CA) as described [23,24]. RNA concentration and quality were determined using a NanoDrop Spectrophotometer (Thermo, MI) and by 1% agarose gel electrophoresis, respectively. Reverse transcription reactions were performed to synthesize cDNA from 200 ng of total RNA using the iScript™ Select cDNA Synthesis Kit (Biorad, CA). Real-time quantitative PCR (qRT-PCR) was carried out in the CFX Connect Real-Time PCR Detection System (Biorad, CA) using iTaq™ universal SYBR^®^ Green supermix (Biorad, CA). The PCR parameters were as previously described [10,19]. Quantification cycle (Cq) values for all genes were compared and analyzed by using the ΔΔC(t) method [25]. All primer pairs used for the analysis of *UCP1, UCP3, TGR5,* and actin were as described [10]. Beta-actin was used as a reference gene.

### High Performance Liquid Chromatography-Electrochemical Detection (HPLC-EC)

The plasma samples collected from each rat were purified and norepinephrine (NE) was extracted according to the combined methods of Denfeld et al [26] and Holmes et al [27]. After extraction, NE levels were assessed using HPLC-EC (Shimadzu, Canby, OR). The mobile phase consisted of 14% methanol, and an 86% mixture of 0.05 M phosphate, 0.03 M citric acid buffer, 0.6 mM octasulfonic acid, and 1.0 mM EDTA-disodium. The pump flow rate was 0.55 ml/min and was set to an operating temperature of 30 °C. NE was separated using a PP-ODS II reverse phase C18-column (Shimadzu, Colombia, MD) and identified according to the retention time of the standard, and concentrations were quantified by comparison with peak heights of the standard concentration curve (10^4^-10^8^ pg/μL). The quantification of sample NE concentrations was performed using an Epsilon electrochemical (EC) detector connected to the HPLC. The detector sensitivity was 5 uA and the oxidation potential was fixed at +700 mV using a glassy carbon working electrode versus an Ag/AgCl reference electrode. Lab Solution software was used to integrate and analyze the raw data for determination of norepinephrine levels.

### 16S rRNA Gene Analysis

Once tolerance to methylone-induced hyperthermia was displayed (4^th^ week of treatment), one day before FMT, daily fecal dropping collection was initiated. This was considered day 0 which was one day prior to the initiation of FMT. During the 7 days of FMT, individual animal fecal droppings were collected, and the cages changed. On the 7^th^ day of FMT, fecal droppings were collected (day 7) followed by treatment with methylone. Collected feces were stored at −70°C.

Isolation of DNA from fecal droppings was performed using a DNeasy Powersoil Kit (Qiagen Inc., CA). The concentration and quality of the DNA were determined using a NanoDrop Spectrophotometer (Thermo, MI) and by 0.8% agarose gel electrophoresis, respectively.

Isolated DNA samples were submitted to LC Sciences (Houston, TX) for standard metagenomic analysis, using primers (341F/805R) that amplify an ~465 bp region containing the V3 and V4 regions of the 16S rRNA gene. The amplified library was sequenced on a NovaSeq platform as paired-end reads (2×250 bp). The resultant raw data were processed by LC Sciences, using a Divisive Amplicon Denoising Algorithm (DADA2)(35), followed by construction of Operational Taxonomic Units (OUT)[24].

### Statistical Analysis

GraphPad InStat v.6.0 software was used to complete all statistical analyses of data except the metagenomic data set. The results are presented as the mean ± SEM of the rectal core body temperatures of the treatment/control groups. Within treatment group changes in body temperature over time were compared with a one-way ANOVA followed by a Dunnet’s post-hoc test. When only two groups were compared, a two-tailed t-test was performed. Significance was established at p<0.05 a priori.

Linear regression and correlation coefficients were determined by plotting individual data points for each subject within the MHT and MN groups (n = 6) for maximal change in temperature following methylone administration and NE levels. The linearity of relationships between plasma NE and maximal change in temperature were determined by linear regression analysis. Statistical significance was determined using a linear relationship ANOVA test.

## Results

### Methylone-induced changes in body temperature

A two-tailed t-test of maximal temperature change comparing MHT and MN controls yielded a significant hyperthermic effect (p = 0.004) on the first week of treatment (Figure 1A). In the second week, significant hyperthermia was again achieved in the MHT group compared to MN controls (p = 0.0001). By week three, hyperthermia was still achieved by MHT rats (p = 0.001) compared to maximal temperature change in the MN group. A one-way ANOVA with Dunnett’s post-hoc test within the MHT group over the five-week treatment period demonstrated no difference in the rise in body temperature between weeks 1 and 2 (p = 0.79); a significantly lower hyperthermic response (p = 0.04) was seen at week three compared week 1. By the fourth week, the hyperthermic response was lost in the MHT group and the temperature did not differ from the MN group (p = 0.881). Throughout the first four weeks of treatment, saline injections did not have an effect on body temperature in the MN group when compared back to the first week of treatment.

**Figure 1.**
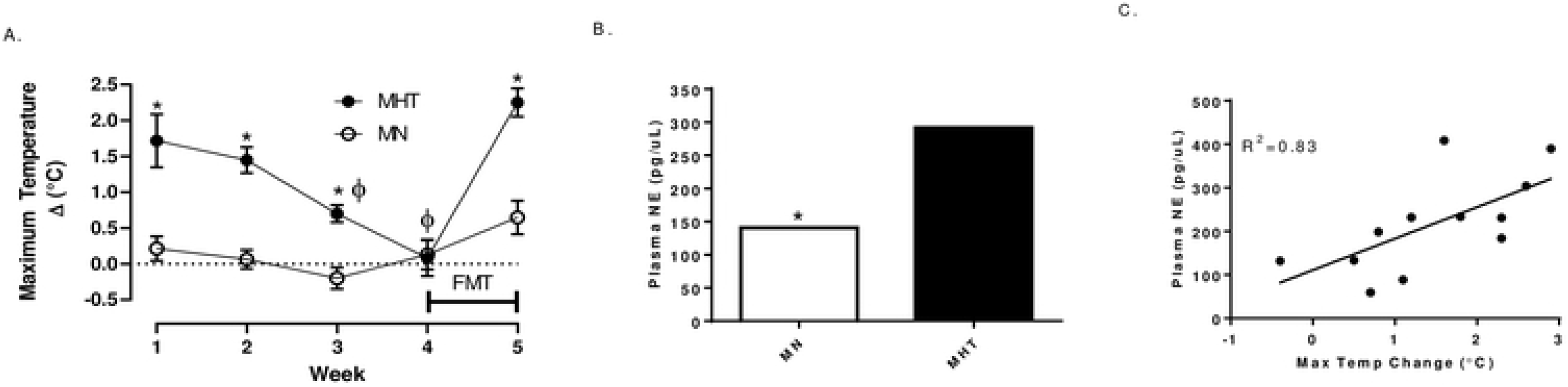
Maximal temperature and norepinephrine (NE) changes associated with once-a-week dosing of methylone (10 mg/kg, sc). (**A**) Weekly maximal temperature changes. “FMT” bar between weeks four and five indicates the duration of the fecal microbial transplant between treatments. *indicates significant difference between each week’s maximal temperature change between the MHT and MN group (p <0.004). ϕ indicates significantly different from MHT maximal temperature change week 1 (p<0.04). (**B**) NE plasma concentrations (pg/μL) for MHT and MN rats following the 5^th^ week of treatment with methylone (10 mg/kg, sc). Each value is the average +/− SEM (n=6). *indicates significant differences from MHT group (p = 0.008). (**C**) Linear correlation between NE levels and maximal temperature changes following methylone treatment. Significance as determined by linear regression ANOVA analysis is shown, along with the correlation coefficients.

Following FMT between weeks four and five, MHT rats exhibited a complete loss of tolerance to methylone-induced hyperthermia, yielding a substantially increased maximal temperature change (p = 0.0001) compared to the previous week where total tolerance was seen. A one-way ANOVA with Dunnett’s post-hoc test comparing week five to week one baseline when the original methylone dose was administered demonstrated that the tolerance effect was lost, and that hyperthermia had returned in the MHT group. The MN group, who received their first dose of methylone at week 5 following the FMT, did not display a significant change in temperature following methylone treatment (p = 0.29).

### Methylone-induced changes in plasma norepinephrine levels

Plasma samples obtained from both treatment groups showed significant increases in circulating norepinephrine levels 90 minutes after methylone (Figure 1B). MN rats that received FMT from MHT rats had lower plasma norepinephrine levels than the corresponding MHT rats that received FMT from the MN animals. Linear regression analysis was performed to compare each animal’s norepinephrine level and the maximal change in temperature induced by methylone. The results demonstrated a significant relationship between plasma levels of norepinephrine and maximal temperature change (Figure 1C).

### Expression of genes involved in methylone stimulation are modulated by FMT

For brown adipose tissues (BAT), qRT-PCR demonstrated a 43-fold increase in the expression of *TGR5* in the MHT group relative to the MN group (Figure 2A). The expression of *UCP1* and *UCP3* (1.48- and 1.38-fold, respectively) was also observed for MHT group relative to that of MN group. Conversely, in skeletal muscle (SKM), both *TGR5* and *UCP3* gene expression was decreased by 5- and 2.5-fold, respectively in MHT group (Figure 2B). Expression of *UCP1* in SKM was below the level of detection.

**Figure 2.**
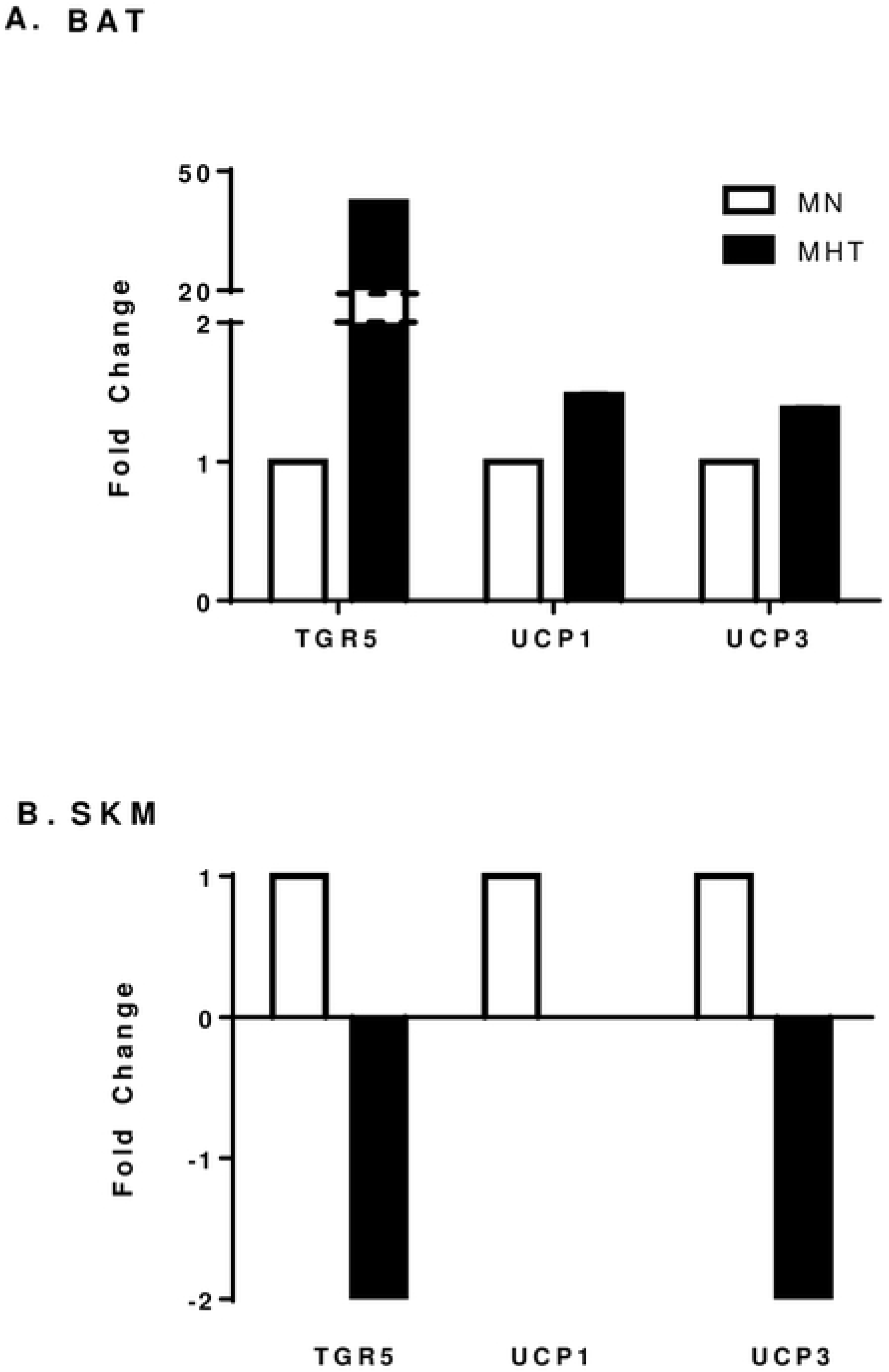
**(A**) Relative fold changes in gene expression of *TGR5*, *UCP*1 and *UCP3* by qRT-PCR of MHT group compared to MN group in brown adipose tissue (BAT) and (**B**) in the skeletal muscle (SKM). Expression of UCP1 in SKM was below the level of detection. Each bar represents the mean ± SEM of three samples performed in triplicate.

### Fecal microbiome composition changes associated with FMT

While fecal samples were collected for all animals before and after FMT, not all samples yielded high quality DNA, as defined by DNA concentration and genome integrity as evidenced by development on 0.8% agarose gels (data not shown). Therefore, we limited microbiome comparisons to six animals (three MHT, and three MN) for which high quality DNA had been isolated both prior to (day 0) and following (day 7) FMT.

Principal coordinates analysis (PCoA) identified MN and MHT animals as having distinct fecal microbiomes, with greater intra-group similarity in the MHT group than in the MN group (Figure 3). Taxonomic comparisons at the Phylum level indicated that for both MN and MHT animals, ~80% of the fecal microbiota taxa consisted of *Firmicutes* (~60%) and *Bacteroidetes* (20%), with the *Proteobacteria* a distant third at 1-6% (Figure 4A). Taxonomic comparisons at the Class level provided greater resolution, with changes in several taxa evident following FMT. MHT recipients demonstrated a marked increase in the relative proportion of the *Firmicutes* Class *Erysipelotrichia*, whereas the reciprocal transplant did not significantly alter *Erysipelotrichia* levels in MN recipients (Fig 4B, Table 1). Notably, the relative proportion of *Erysipelotrichia* was four-fold less in the MHT group than in the MN group. Conversely, for the *Proteobacteria* Classes *Gammaproteobacter* and *Alphaproteobacter,* relative proportions were similar between MHT and MN animals prior to transplant but increased several folds in each of the MN animal recipients following FMT from MHT donors. In the reciprocal transplants, the relative abundance of *Gammaproteobacter* remained similar for two of the three MHT recipients, with the third recipient showing a multifold increase in the relative abundance of *Gammaproteobacter* (Fig 4B, MHT6, Table 1). Similarly, *Alphaproteobacter* displayed multi-fold increases in each of the three MN animal recipients following FMT, and little change for two of the three MHT recipients, again with same animal showing a several-fold increase following FMT from the MN donor (Fig 4B, MHT6; Table 1).

**Table 1.**
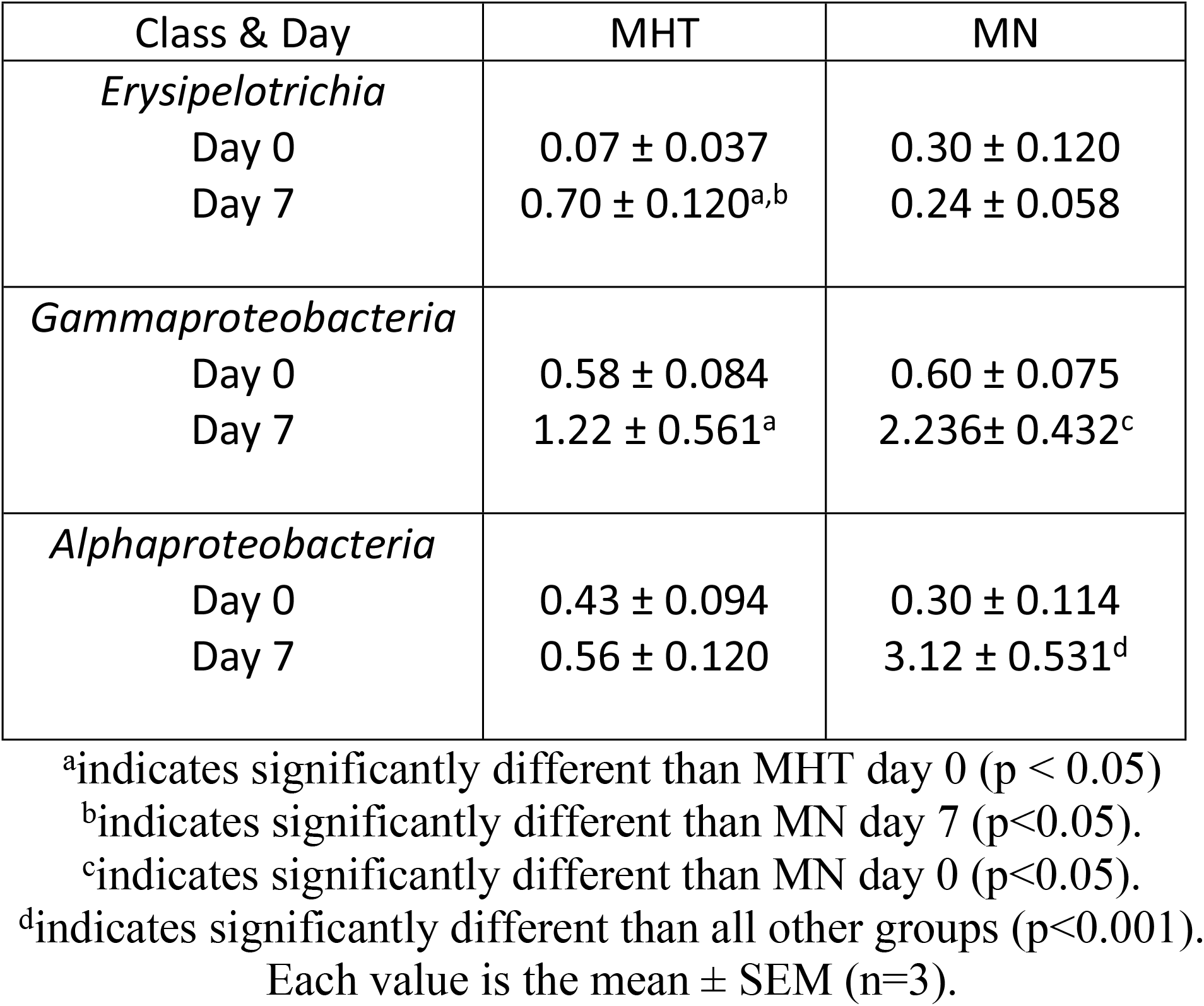
Mean relative percentage of microbiotia as the Class *Erysipelotrichia*, *Gammaproteobacteria* and *Alphaproteobacteria* pre-(day 0) and post-(day 7) FMT

| Class & Day | MHT | MN |
| --- | --- | --- |
| *Erysipelotrichia* |  |  |
| Day 0 | 0.07 ± 0.037 | 0.30 ± 0.120 |
| Day 7 | 0.70 ± 0.120<sup>a,b</sup> | 0.24 ± 0.058 |
| *Gammaproteobacteria* |  |  |
| Day 0 | 0.58 ± 0.084 | 0.60 ± 0.075 |
| Day 7 | 1.22 ± 0.561<sup>a</sup> | 2.236 ± 0.432<sup>c</sup> |
| *Alphaproteobacteria* |  |  |
| Day 0 | 0.43 ± 0.094 | 0.30 ± 0.114 |
| Day 7 | 0.56 ± 0.120 | 3.12 ± 0.531<sup>d</sup> |

<sup>a</sup>indicates significantly different than MHT day 0 (p < 0.05)
<sup>b</sup>indicates significantly different than MN day 7 (p<0.05).
<sup>c</sup>indicates significantly different than MN day 0 (p<0.05).
<sup>d</sup>indicates significantly different than all other groups (p<0.001).
Each value is the mean ± SEM (n=3).

**Figure 3.**
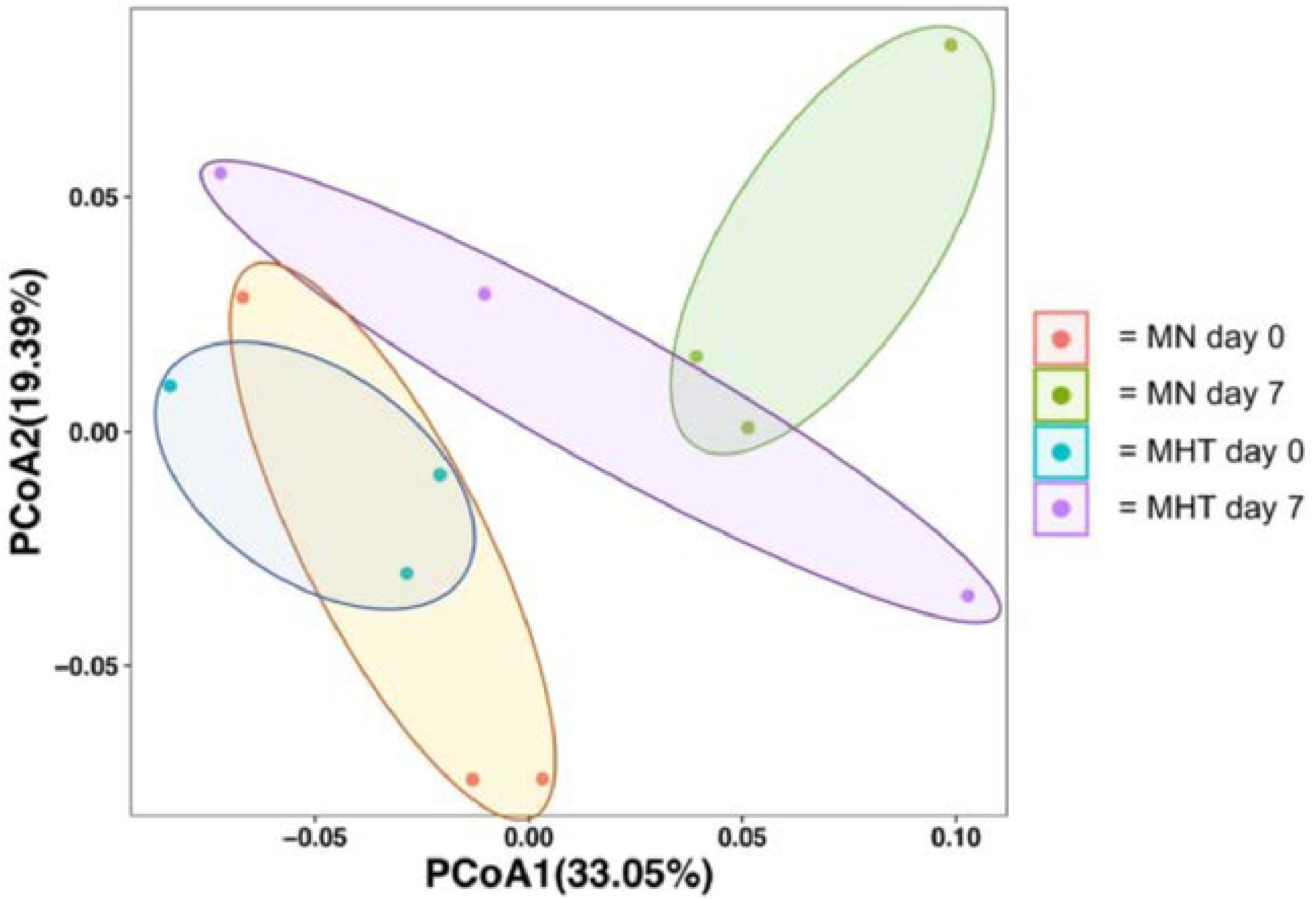
Principal coordinate analysis (weighted, unfractionated) of overall microbiome composition pre- and post-FMT. Colored circles represent the microbiome of an individual at the 95% confidence interval, with the group membership identified by color, as indicated in the key, where; “MN day 0” (orange circles) represents untreated control animals prior to FMT, “MN day 7” (green circles) represents untreated control animals following to FMT, “MHT day 0” (blue circles) represents methylone tolerant animals prior to FMT and “MHT day 7” (purple circles) represents methylone tolerant animals following to FMT. p = 0.019.

**Figure 4.**
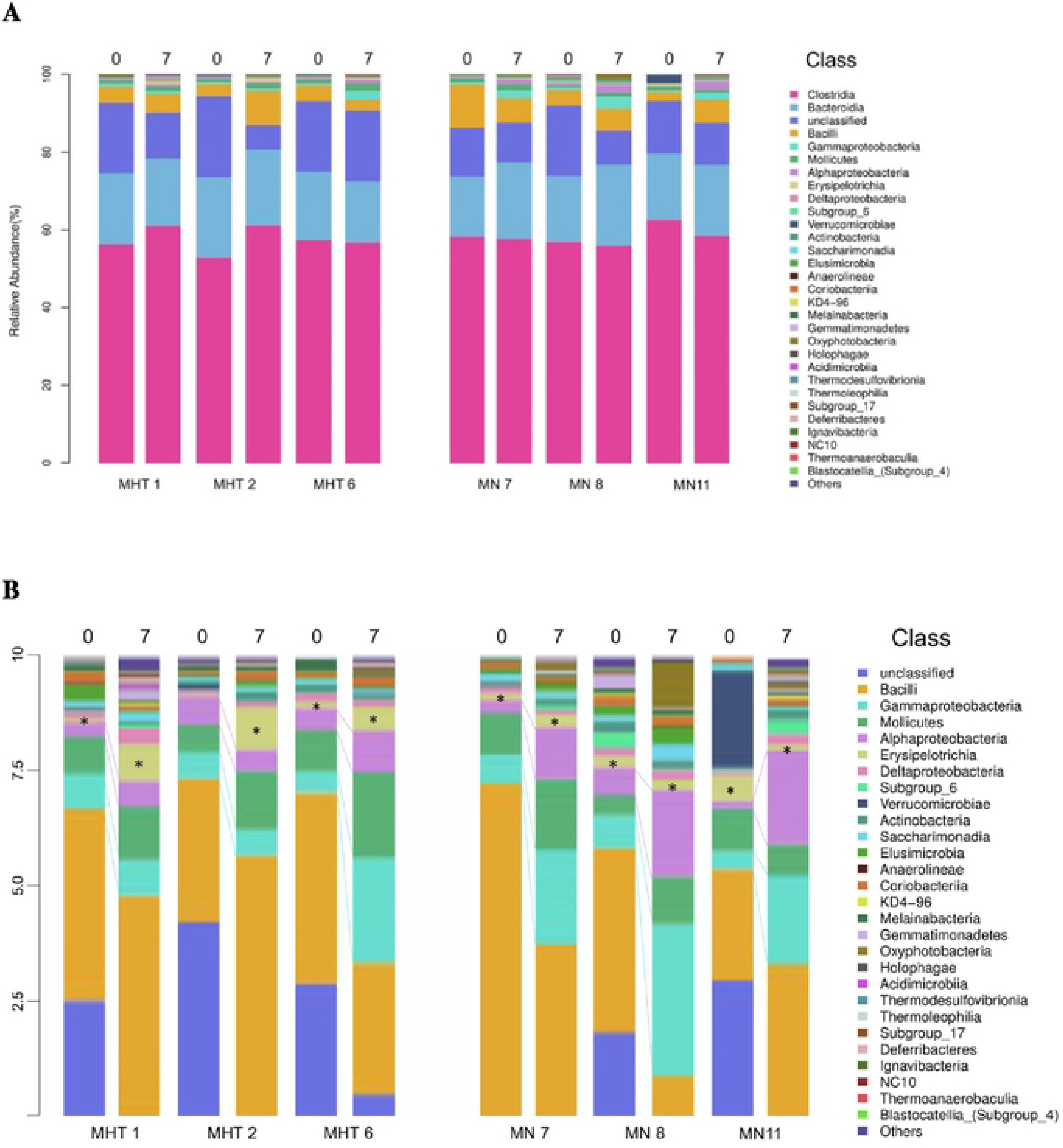
(**A**) The relative abundance of the 30 most common bacterial Classes identified for individual animals prior to (“0”) and following (“7”) FMT are depicted as stacked bars, with individual Classes identified by color, as indicated by the key at the right. Methylone hyperthermia-tolerant (MHT) and methylone-naive (MN) animals are identified by number. (**B**) The upper 10% of Fig 5b; vertically stretched 10x to allow comparison of less prevalent Classes pre- and post-FMT. Changes in the relative number of *Gammaproteobacteria* (light blue) and *Alphaproteobacteria* (light purple) are highlighted by dotted lines between the pre-FMT (“0”) and post-FMT (“7”) stacks for each animal. Boxes corresponding to *Erysipelotrichia* (pale yellow) are highlighted as “*”.

## Discussion

Here, we demonstrate for the first time that FMT can influence the temperature response changes induced by sympathomimetic agents such as methylone. By the fourth week of treatment, tolerance to the drug-induced hyperthermia had developed. Following FMT between weeks four and five, MHT rats exhibited a complete loss of tolerance to methylone-induced hyperthermia, yielding a substantially increased maximal temperature change compared to the previous week where total tolerance was observed. Moreover, the MN group who received their first methylone treatment and would normally be expected to experience significant increases in body temperature, displayed no significant change in temperature after receiving the FMT from MHT donors. These findings are consistent with a previous study that suggested the potential link between sympathomimetic-induced thermogenesis and gut bacteria [10], and strikingly, provided evidence that the FMT is capable of modulating the thermogenic response. To our knowledge, these results appear to be the first evidence to demonstrate that a pharmacologically mediated temperature response to a drug can be reproduced from donor to recipient through fecal microbial transplant in an animal model. In this case, based on the norepinephrine differences between the treatment groups, FMT had a substantial influence on the sympathetic nervous system and introduced a tolerance effect to the drug naïve group that would otherwise only occur in subjects that had been repeatedly exposed to the drug.

The restoration of methylone-induced hyperthermia in the MHT group was associated with a significant increase in plasma norepinephrine levels as compared to the MN group which did not display a hyperthermic response. The gut-microbiota has been shown to produce catecholamines [28], and norepinephrine has been suggested to play a key communication role in the microbiome-gut-brain axis. Additionally, exogenously administered norepinephrine can induce *Escherichia coli* chemotaxis, motility, and virulent gene expression [29] through binding to the bacterial norepinephrine-like receptor, QseC [30]. Given that sympathomimetic agents can increase plasma norepinephrine up to 35-fold [31], it is not surprising that these agents can influence the gut microbiome and vice versa.

UCP1 and UCP3 have further been demonstrated to play complementary roles in the onset (UCP1) and maintenance (UCP3) of sympathomimetic-induced hyperthermia [32]. We have previously demonstrated that in male rats, tolerance to methylone-induced hyperthermia occurs between the fourth and fifth weeks following once a week treatment [19]. In that study, the gene expression levels for *UCP1*, *UCP3* and *TGR5* were also measured in brown adipose tissue (BAT) and skeletal muscle (SKM). Tolerance was associated with an increase in *UCP3* in BAT and increases in *UCP1* and *UCP3* in skeletal muscle [19]. Here, following FMT, BAT demonstrated an increase in the expression of *TGR5, UCP1, and UCP3* in the MHT group relative to the MN group. Conversely, in SKM, both *TGR5* and *UCP3* gene expression was decreased in the MHT group. These changes are consistent with the key roles UCPs play in mediating sympathomimetic hyperthermia and the restoration of the hyperthermic response following FMT in the MHT treatment group.

Previously, a hyperthermic dose of MDMA was shown to lead to the enrichment of the relative proportion of a *Proteus mirabilis* strain in the ceca [10]. In that same study, antibiotic treatment not only prevented the *Proteus mirabilis* enrichment but also attenuated MDMA-induced hyperthermia. Angoa-Perez et al., [33] examined the effects of synthetic cathinones on the diversity and taxonomic structure of the gut microbiome. Those authors found that the two phyla most altered by the synthetic cathinones were *Firmicutes* and *Bacteriodetes*. Similarly, in the present study, taxonomic comparisons at the Phylum level indicated that for both MN and MHT animals the greatest effects were also on *Firmicutes* and *Bacteroidetes*. As noted by Angoa-Perez et al., [33], this effect on *Firmicutes* and *Bacteroidetes* is expected as these two phyla are dominant in rodents [33]. The specific identity of the microbe(s) involved in the present temperature changes is unknown due to insufficient resolving power of 16S rRNA gene alone; however, the concordance of changes in the relative abundance of *Gammaproteobacter* and *Alphaproteobacter* following FMT implicates members of these two phyla as potential contributors to the establishment of methylone tolerance. The lower relative abundance of *Erysipelotrichia* in MHT animals, and its increase following FMT similarly implicates this phylum as a potential contributor.

There are a number of critical considerations to be made in the interpretation of these data. While the roles that clinical and recreational agents have in contributing to dysbiosis of the microbiome have just begun to be characterized and the overall effects appear to be compound specific, there is often significant individual variation between microbe communities in test subjects [34, 35], complicating the interpretation of the changes induced by the drugs. Based upon our experiments, we do not know whether methylone itself is directly mediating changes to the microbiome or if these changes are indirect and secondary to a pharmacodynamic response (e.g., hyperthermia) to the drug. Although the findings in the present study suggest a link between the gut-brain axis and sympathomimetic-induced hyperthermia, the gut microbiome changes may also be playing a peripheral role in altering the thermogenesis mediated by methylone. Finally, the use of FMT may have selected for aerobic or facultative anaerobic taxa which may be reflected in our post-FMT taxonomic differences.

## Conclusion

The bi-directional FMT between MHT and MN resulted in a complete reversal of the predicted hyperthermic response in the MN group. After displaying tolerance to the hyperthermia mediated by methylone, the FMT from MN to MHT resulted in a return of hyperthermia in animals that over a four-week treatment period continued to display resistance. Given that the gut microbiota has been demonstrated to impact thermoregulation in general [12], the results from the present study further support the contention that the gut microbiome plays a contributing role in the hyperthermia mediated by sympathomimetic agents such as methylone, and that fecal microbial transplants may be able to transfer phenotypic responses to pharmacological agents.

